# Encoding innate ability through a genomic bottleneck

**DOI:** 10.1101/2021.03.16.435261

**Authors:** Alexei Koulakov, Sergey Shuvaev, Divyansha Lachi, Anthony Zador

## Abstract

Animals are born with extensive innate behavioral capabilities, which arise from neural circuits encoded in the genome. However, the information capacity of the genome is orders of magnitude smaller than that needed to specify the connectivity of an arbitrary brain circuit, indicating that the rules encoding circuit formation must fit through a “genomic bottleneck” as they pass from one generation to the next. Here we formulate the problem of innate behavioral capacity in the context of artificial neural networks in terms of lossy compression of the weight matrix. We find that several standard network architectures can be compressed by several orders of magnitude, yielding pre-training performance that can approach that of the fully-trained network. Interestingly, for complex but not for simple test problems, the genomic bottleneck algorithm also captures essential features of the circuit, leading to enhanced transfer learning to novel tasks and datasets. Our results suggest that compressing a neural circuit through the genomic bottleneck serves as a regularizer, enabling evolution to select simple circuits that can be readily adapted to important real-world tasks. The genomic bottleneck also suggests how innate priors can complement conventional approaches to learning in designing algorithms for artificial intelligence.

## Introduction

Many animals are born with impressive and elaborate behavioral capacities. Soon after birth, a spider can build a web, a whale can swim, and a monkey fears snakes. From an evolutionary perspective, it is easy to see why such innate abilities would be selected for: Those individuals that can survive beyond their most vulnerable early hours, days, or weeks are more likely to survive until reproductive age and hence produce progeny at a higher rate. Of course, in practice there is no crisp distinction between innate and learned abilities; innate abilities form a foundation for learning, and animal behavior arises from an interaction between these two processes. Although learning has been studied extensively in the context of artificial intelligence, there has been much less theoretical attention devoted to the structure of innate behaviors.

Innate behaviors are encoded in the genome and can be expressed in the neural circuits already present at birth. However, this poses a challenge: How can a complex neuronal connectivity diagram be encoded into a genome? The size of the genome provides an approximate upper bound on the amount of information transmitted from generation to generation. The genome of the simple worm *C. elegans* is about 10^8^ base pairs (Bargmann, 1998), so it could transmit up to about 2 × 10^8^ bits. This in principle would be more than adequate to explicitly encode the highly stereotyped connectivity among the 302 neurons in the *C. elegans* brain, since even a dense 302^2^ connection matrix would take at most 9 × 10^4^ bits to store, times a small factor associated with the number of bits per synaptic weight. On the other hand, the human genome is only about an order of magnitude larger than that of *C. elegans* (∼10^9^ bits), whereas the human cortex has about 10^10^ neurons, so even the (sparse) cortical connectivity matrix might require at least 10^15^ bits to specify. This implies that the human cortex would require about 5–6 orders of magnitude more information to specify than is available in the genome, if every connection were specified explicitly (Zador, 2019). Since the genome encodes the rules for wiring up the nervous system, a natural question is how the small amount of information contained in the genome can instruct the creation of the large capacity cortex. We refer to the long recognized (Tessier-Lavigne and Goodman, 1996; Sperry, 1963; Stanley et al., 2019) mismatch between the information capacity of the genome and the complexity of the resulting neural circuit as a “genomic bottleneck” (Zador, 2019).

The mismatch described by genomic bottleneck implies that the connectivity of most neuronal circuits, including the mammalian cortex, is not explicitly specified, neuron-by-neuron, in the genome. Rather, the genome specifies rules for connectivity. It has long been recognized that simple rules can give rise to surprisingly complex structures (Turing, 1952; Von Neumann et al., 1951), and it is straightforward to formulate simple, low-complexity rules that specify the connectivity in arbitrarily large networks. For example, the simple rule “connect to your four nearest neighbors” specifies a grid of potentially unlimited size (Fig. 1A). Another class of rules specifies connections between cells based on the surface markers they express (Sperry, 1963; Zipursky and Sanes, 2010; Wei et al., 2013) (Fig. 1B); axons can exploit these markers to find their destinations (Goodhill and Baier, 1998). The columnar organization that is observed in many brain regions allows for the replication of similar connectivity modules throughout the brain, thus limiting the number of parameters needed to wire the circuit (Itzkovitz et al., 2008). Complex structures such as orientation columns in the visual cortex can be induced to self-organize from simple use-dependent rules (Von der Malsburg, 1973). Developmental rules such as these can dramatically reduce the amount of information needed to specify the connectivity of a neural circuit, prior to extensive experience.

**Figure 1:**
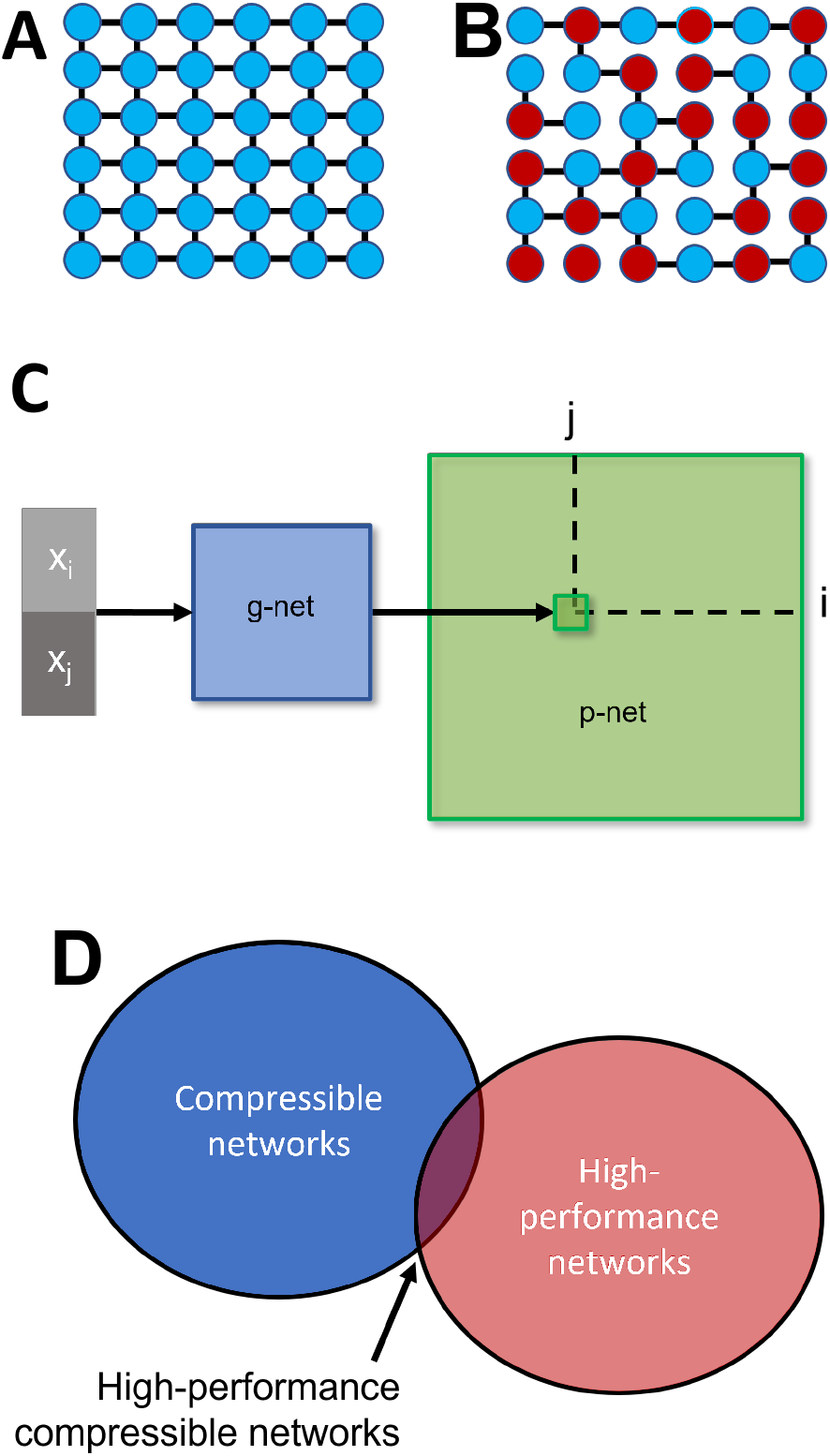
Simple rules can specify networks. (A) A very simple nearest neighbor wiring rule. (B) A somewhat more complex rule (only connect to nearest neighbors of opposite color) leads to a more complex network. (C) Network specification through a genomic bottleneck. The input to the genomic network (“g-network”) is a pair of neurons (pre- and postsynaptic) specified by binary strings. Each neuron has a unique label, consisting of unique binary string. The two binary labels are passed through the g-network, which assigns the strength of the connection between the two neurons in the “p-network.” Because the number of parameters in the g-network is smaller than the number of parameters in the p-network, the g-network is effectively compressing the p-network. (D) G-networks seek to discover p-networks that both solve the problem well and are compressible.

In spite of this extensive literature on modeling development, these simple developmental rules do not typically specify networks with the capacity to perform complex general computations. Thus, although such rules can readily specify the formation of repeated modules that enable the emergence of receptive fields in the retina or the visual cortex (Linsker, 1986), such stereotyped modules cannot directly encode more specialized knowledge like a spider’s capacity to build a web or a rat’s innate fear of fox odor (LeDoux, 2012). We therefore set out to explore how low-complexity rules can give rise to networks that perform complex well-defined computations. Within the framework of artificial neural networks (ANNs), we seek to compress the complex connectivity (weight matrix) into a much smaller “genome.” The decoding of this genome into the initial weights of the network enables the network to perform well upon initialization, without additional training. This decoding is analogous to the neurodevelopmental processes by which the genome provides a blueprint for circuits that enable animals to perform essential tasks at or soon after birth. We hypothesized that under some conditions, compressing the weight matrix through a “genomic bottleneck” would extract the most useful and important features of the connectivity; the genome would act as an “information bottleneck” (Tishby et al., 2000; Saxe et al., 2019; Tishby and Zaslavsky, 2015). In this way, a physical constraint—the limited size of the genome—might actually be an algorithmic advantage, serving as a regularizer and thereby turning a potential “bug” into a feature.

### Implementation of the Genomic Bottleneck

To test these ideas, we first trained standard feedforward ANNs on well-studied supervised learning tasks. ANNs consist of nodes (“neurons”) connected by weights (“synapses”). Artificial neural networks are typically initiated randomly—*tabula rasa*—and acquire their functionality through learning. When an ANN learns a task, the “knowledge” of the task is summarized in the weights of the ANN. We use “connectivity” to describe both the specification of which connections are non-zero, as well as the strengths or weights of those connections. We refer to the trained network as the “phenotype network,” or “p-network”.

We sought to compress the p-network through a genomic bottleneck, preserving as much of the performance as possible. The compressed network serves to initialize the network, endowing it with innate abilities prior to learning. To search widely over the space of possible compressions, we used a separate ANN—a “genomic network” or “g-network”—to generate the p-network (Fig. 1C). The formulation of the neurodevelopmental process as an ANN allows us to focus on the genomic bottleneck at a conceptual level, without the need to model the complexities of neural development. This leads to a model in which genomes, and the circuits they encode, are co-optimized in nested loops: an inner loop corresponding to “learning” in animals, and an outer loop corresponding to “evolution.” For reasons of efficiency, in our model both the inner and outer loops are optimized by gradient descent, and are not intended as detailed models of learning or evolution.

The inputs to the g-network are a pre- and postsynaptic pair of neurons, each represented by a unique binary vector; and the output of the g-network is the expected strength of the connection between these neurons. For sparsely connected networks, the strength will often be zero. This formulation is inspired by neurodevelopmental rules based on local pair-wise interactions between neurons (Sperry, 1963; Zipursky and Sanes, 2010), but it is not intended as a realistic model of neural development (Stanley et al., 2009; Barabási and Barabási, 2020). To facilitate the efficient search for g-networks that achieve good compression, we used stochastic gradient descent (Ha et al., 2016) rather than evolutionary algorithms (Stanley et al., 2009, 2019) to achieve end-to-end optimization of both the g-network and the p-network.

We represented each neuron in the p-network by a unique binary vector. Each binary digit in this label can be interpreted as an indicator of the presence of one type of a “molecular tag” in the expression profile of this neuron. Thus, for example, if each neuron is represented by 10 binary digits, the input layer will consist of 20 units (10 each for presynaptic and postsynaptic neurons). Connectivity is effectively guided by interactions of pairs of molecules expressed on the pre- and postsynaptic membranes. The g-network framework can naturally accommodate the addition of new neurons, new layers, and new connections between layers. A network with *N* neurons requires *log*_2_(*N*) binary tags to specify each neuron uniquely, so the input to the g-network is *log*_2_(*N*) units (binary digits) for each neuron. Because the size of the g-network’s input grows only logarithmically as more neurons are added to the p-network, the g-network can implement connectivity of arbitrarily large p-networks without substantial changes in size. Thus, the small amount of information parameterizing the g-network can be used to encode even very large p-networks.

Neuronal projections are often organized topographically. For example, nearby ganglion neurons in the retina encoding nearby points in physical space project topographically via the thalamus to nearby points in visual area V1. However, we did not explicitly encode geometric space into the network (Stanley et al., 2009). Instead, we adopted a more general approach in which each neuron was labeled according to a two-dimensional Gray code (Frank, 1953), so that the binary vectors representing nearby neurons differ by only a small number of bits (Supp Fig. 1A). The Gray code facilitates the discovery of connectivity rules that exploit spatial structure, but also has the potential to allow the network to discover wiring motifs associated with distinct neuronal “types” or “classes”. For example, one bit of the neuronal specification in the Gray’s code could represent the distinction between excitatory and inhibitory neurons, and another bit could denote a particular subtype of inhibitory neuron. In this way, the characteristic connectivity patterns associated with specific neuronal subtypes in brains (Kepecs and Fishell, 2014) could readily arise in these ANNs.

Although a sufficiently large g-network could, in principle, perfectly recapitulate the p-network by “memorizing” all of the connections exactly, this would fail to compress the p-network, and thus would be unlikely to extract more general wiring motifs. We therefore focused on a regime where the size (complexity) of the g-network is substantially smaller than the size of the p-network, encouraging the g-network to discover compact wiring rules. Our goal is to discover network architectures that are both high performance and compressible (Fig. 1D). This setting can be viewed as a lossy compression problem, where the success of compression is measured not by, e.g., the reconstruction error as in typical lossy image compression, but rather by the ability of the uncompressed weight matrix to perform well on the target task without further training.

Our approach can be described as the search for a solution that balances two competing goals. First, we wish to achieve good innate performance, i.e. we wish to minimize the error *E*_0_(*G*), where *E*_0_(*G*) is the error before training of the p-network encoded by the corresponding g-network *G*. Second, we also wish to limit the complexity of the genomic network, specifically the entropy *H*(*G*) of the genome *G*. We use the number of parameters used to specify the g-networks as a surrogate for the entropy *H*(*G*). Conceptually, we can thus formulate our overall goal as a lossy compression problem in which we seek to minimize an objective function *J* with respect to the genome parameters *G*:

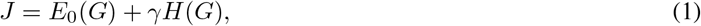

where *γ* is a positive parameter that specifies the tradeoff between the two goals. (In practice, we select a particular complexity *H*(*G*) and then minimize the error for that network). In this formulation, the second term can be seen as a regularizer, related to techniques such as weight pruning (LeCun et al., 1990; Han et al., 2015), that seek to keep the weight matrix simple.

We used an iterative algorithm to find g-networks which generate p-networks that are both compressible and achieve good ab initio performance (see Methods). First, we used standard stochastic gradient descent to find a weight matrix *W*^(1)^ of a p-network that minimizes the training error. The elements of this weight matrix are then used to train the weights of a g-network. Specifically, the training set for the g-network consists of inputs {*b*_*i*_, *b*_*j*_} and target outputs 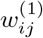, where *b*_*i*_ is the binary representation of the *i*^*th*^ unit of the p-network, and 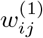 is the connection from unit *i* to unit *j*. The g-network trained in this way thus generates an approximation 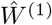 of the p-network weight matrix on which it was trained. This approximation is then used as the initialization to train the next iteration *W*^(2)^ of the weight matrix, and the process is iterated to yield 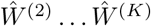. This procedure yields networks that are both capable of accurate initial (innate) performance and occupy limited amounts of space in the genome (compressible).

### Supervised Learning

#### Solving MNIST

We first tested our approach on a classic supervised learning problem: handwritten digit recognition. In this problem, a network is trained with examples of handwritten digits (0-9) taken from the MNIST dataset (Fig. 2A; *see Methods*) and learns to assign labels to new examples of these digits it has not previously encountered. For the p-network we used a standard fully-connected network architecture with 28 × 28 = 784 pixel units at the input layer and one hidden layer with 800 units, for a total of 6 × 10^5^ parameters. With random initial weights, the performance rose from random (10%) to about 98% (Fig. 2C) after about 20 epochs.

**Figure 2:**
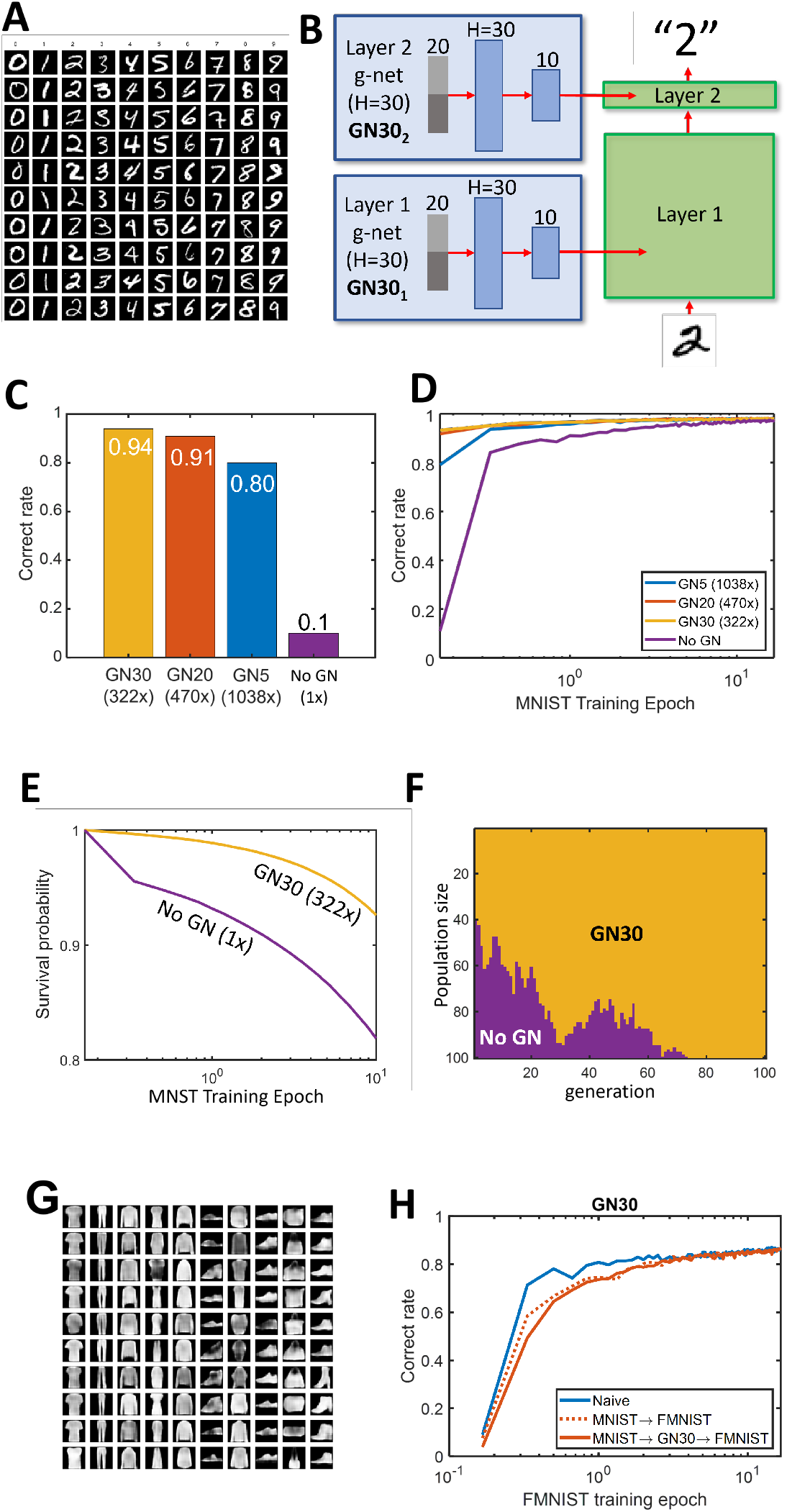
Genomic bottleneck approach to MNIST. (A) Examples of MNIST dataset of handwritten digits. (B) We used a two layer fully connected MNIST network created by individual g-networks for each layer (*GN* 30_1_ denotes a g-network with *H* = 30 units in the hidden layer for the second MNIST layer). GN30 corresponds to 322-fold compression. (C) Initial performance for several levels of compression. Performance is excellent even with 1038-fold compression. (D) Learning dynamics for different levels of compression. (E) Fitness advantage of model organism with high innate performance. (F) Model organism with high innate performance dominates population. (G) Example of fashion-MNIST dataset. (H) Failure of transfer learning f-MNIST dataset. Blue, dotted and solid red lines represent results for training a network using FMNIST dataset that is initialized by random, MNIST weights, and weights generated using GN30 trained on MNIST data respectively

We then used a much smaller g-network with only 30 hidden units (GN30), or about 2 × 10^3^ parameters, to compress the p-network. The g-network trained using the algorithm described above generated a p-network with 94% correct performance upon initialization. Thus the g-network was able to achieve 322-fold compression, while maintaining innate performance almost equal to that of the fully trained network. We observed a tradeoff between the degree of compression and the innate performance (Fig. 2C,D), but good innate performance of 79% correct could even be achieved using GN5, a network with 1038-fold compression. These results demonstrate that there exist p-networks that are both compressible and high-performance.

To illustrate how high innate performance could provide an evolutionary advantage, we formulated a simple model in which an organism’s survival to reproductive maturity—its fitness—was proportional to its performance on this task. In this model, the probability *p*(*t*) that the organism is still alive at time *t* is given by

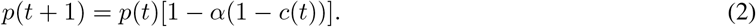

Here, *c*(*t*) is the rate of correct performance at time *t* and 0 < α < 1 is a parameter that determines the contribution of this trait to survival. This model thus relates correct performance on the task to survival (Fig. 2E). Fig. 2F shows how, over successive generations, the fraction of individuals with high innate performance increases at the expense of individuals which rely solely on learning to acquire fitness. As expected, over several dozen generations the individuals with higher innate fitness dominate the population, completely supplanting those initialized *tabulasa*, which must learn everything from the environment. Many factors in the real world could serve to complicate this simple model, which for example does not explain the prolonged period of posnatal helplessness of mammals. Nonetheless, the model provides an intuition for why evolution might be expected to maximize high innate performance.

We hypothesized that passing the wiring diagram through the genomic bottleneck would extract the most useful and important features of the connectivity and enable generalization to related tasks. To test this idea, we used the related “fashion” MNIST, or F-MNIST dataset, which has the same format as the MNIST dataset but consists of ten different categories of clothing (shirts, shoes, etc; Fig. 2F). Disappointingly, we observed no enhancement of F-MNIST learning upon initializing weights using MNIST-trained g-network. Indeed, the p-network adapted from MNIST actually showed somewhat slower learning than a naive network (Fig. 2G), as though the network first had to unlearn MNIST before learning F-MNIST. We hypothesized that this failure to generalize across visual recognition tasks was due to overfitting on the specifics of MNIST dataset, due to the relative simplicity of the tasks and the network used to solve them: Both of these datasets are too simple to require learning general properties of images that can be transferred to novel visual problems.

#### Solving CIFAR10

To test whether there are conditions where the genomic bottleneck might extract features that generalize over multiple datasets, we applied the algorithm to a more complex problem that requires a deeper network. We used the CIFAR10 dataset, which consists of 60k color images drawn from 10 categories such as airplanes, cars, and cats (Fig. 3A). For the p-network we used a standard 9-layer convolutional neural network (CNN) architecture with about 1.4 × 10^6^ weights (*see Methods*). We compressed each CNN layer of the p-network with a separate g-network (Fig. 3B). Similarly to MNIST, the compressed CIFAR10 network reached high performance. GN50, a network with 92-fold compression, achieved initial performance of 76% (vs. naive 10%), fairly close to the fully trained performance of 89% (Fig. 3C, D). Thus, as with MNIST, the g-network achieved approximately two orders of magnitude compression while maintaining good initial performance.

**Figure 3:**
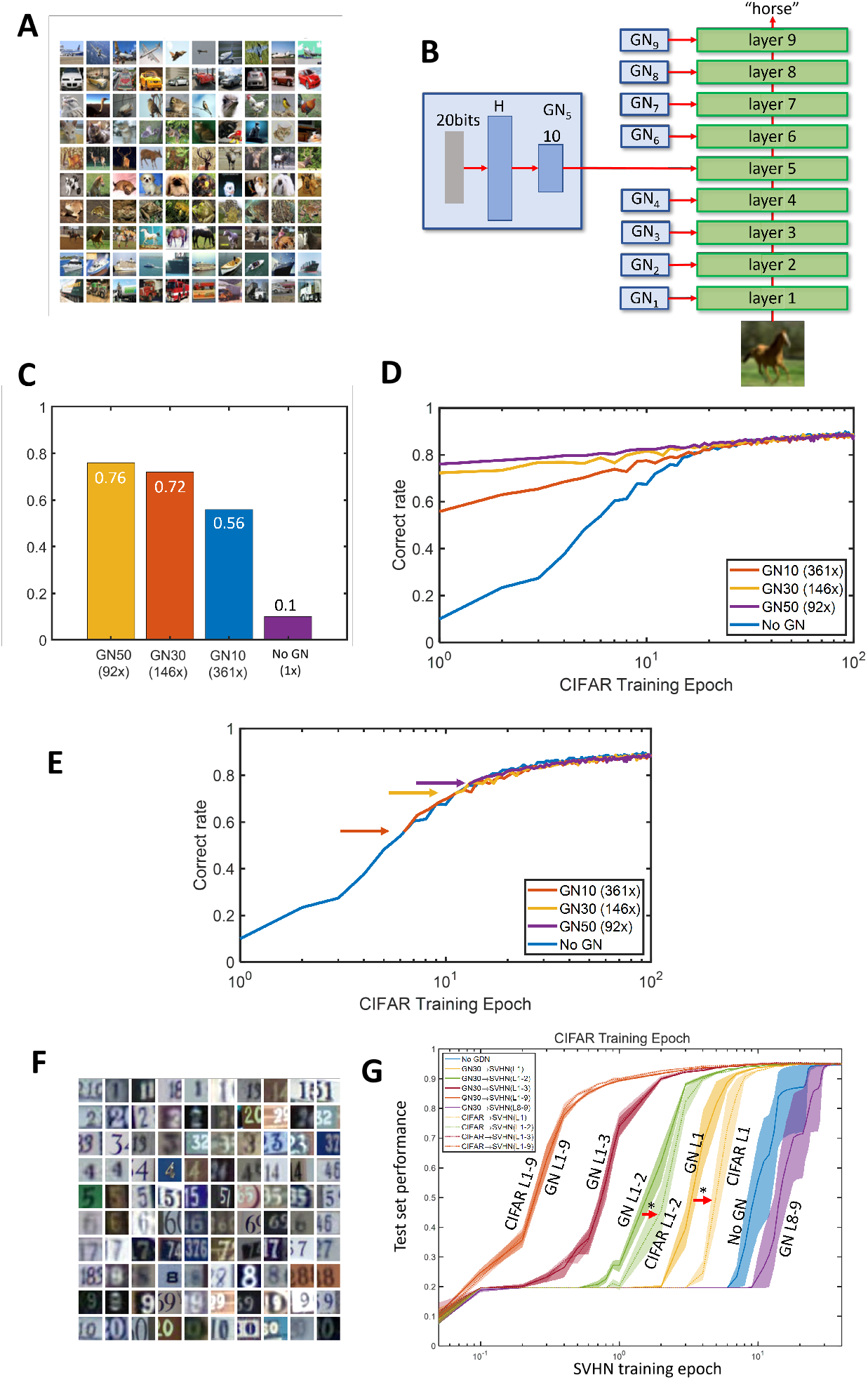
Genomic bottleneck approach to CIFAR10. (A) Examples of CIFAR10 dataset of images. (B) To classify CIFAR10 images, we used a nine layer convolutional network. Each layer was created by individual g-networks. (C) The p-network achieves excellent *tabula rasa* performance even greater than 100-fold compression. (D) Dynamics of learning for different levels of compression. (E) The learning rate of the compressed networks is the same as *tabula rasa* networks. For each level of compression, the curve in D is shifted to the *tabula rasa* curve. (F) Example of SVHN dataset. (G) Transfer learning to SVHN dataset. Results are shown as training curves on SVHN dataset for networks initialized using different sets of layers transferred from CIFAR10 dataset as indicated. For example, green solid curve (GN L1-2) shows results for layers 1 and 2 initialized using g-nets trained on CIFAR10 data, while the remaining layers are initialized randomly (shaded regions show standard deviation of the *mean*). For CIFAR L1-2 curve, layers 1 and 2 were initialized by direct transfer from CIFAR10 dataset. That GN L1-2 is shifted compared CIFAR L1-2 (red arrow) indicates the advantage of our approach. The similar feature is observed for layer one transfer curves (orange). Stars indicate statistical significance (p<0.05). GN L8-9 curve shows worse performance than naive training (No GN), similarly to Fig. 2H.

The learning dynamics (Fig. 3D) demonstrate the utility of genomic compression for achieving enhanced initial performance. However, it was not clear whether this was also associated with faster learning. To our surprise, genomic compression had no effect on the learning trajectory; the only speedup was due to the higher initial performance, as though the p-network was “hot-started” by the g-network (Fig. 3E). Thus genomic compression, at least under these conditions, did not affect the learning rate.

To assess whether the structure that g-networks extracted from the CIFAR10 dataset could be useful for other datasets, we tested transfer from the CIFAR10-trained network to a related problem. We used the Street View House Numbers (SVHN) dataset, which contains images of street numbers in a format similar to the CIFAR10 dataset (Fig. 3F). We first confirmed the effectiveness of a standard algorithm for transfer learning. We trained the p-network on the CIFAR10 dataset without compression, and then used the p-network’s weights as a starting point for SVHN training (CIFAR10 transfer). As expected, this procedure accelerated learning, reducing training time from about 20 training epochs (Fig. 3G, blue line, “No GN”) to a single epoch (Fig. 3G, “CIFAR10 L1-9”). This implies that the CIFAR10 dataset contains features that are similar to SVHN dataset.

We next compared the standard transfer learning algorithm to an algorithm based on the genomic bottleneck. To achieve transfer with the genomic bottleneck, we used g-networks to generate p-network as described above and then used this p-network as an initial condition for SVHN training (g-network mediated transfer). Remarkably, the performance of g-network mediated transfer was indistinguishable from the standard approach (Fig. 3F, orange solid line, “L1-9”), even though in this case the number of transferred parameters was 92 times fewer. These results indicate that whatever is crucial for transfer from CIFAR10 to SVHN is captured by the nearly 92-fold smaller g-network.

To further dissect the consequences of genomic compression, we examined the effect of transferring only one or a few layers at a time. When only the first two lower layers (layers 1-2) were transferred from CIFAR to SVHN, while randomly initializing the remaining layers, genomic transfer actually yielded faster learning than direct transfer (Fig. 3F, red arrows). For example, when layer 1 was initialized with g-network, 50% performance was reached in about 3.2 training epochs, while similar levels were only achieved in 4.8 epochs using direct CIFAR10 transfer—a 1.5-fold difference. Thus, it appears that the lower layers of the network contain features that generalize across datasets, and these features are extracted particularly well using the g-network based compression algorithm. On the other hand, transferring the last two layers of the network resulted in slower training compared to the naive case, a result reminiscent of our previous result with MNIST-to-F-MNIST transfer (Fig. 2H). This result implies that the last two layers of CIFAR10-trained network contained features that are specific to the dataset and were not useful for the recognition of the house numbers in SVHN data.

Taken together, these findings demonstrate that g-networks can extract structure that is generalizable across datasets. Compression with g-networks yields performance that is comparable to—and in some cases better—than simple uncompressed weight transfer, indicating that g-networks identify a special subclass of p-networks that are compressible and capture essential structure of the data (Fig. 1D). This enhancement is particularly evident in the rate of transfer of the lower layers in deep nets (Fig. 3G). Interestingly, the receptive fields of neurons in the lower visual system show substantial similarities between different species, while higher layers are more specialized (Rodieck and Rodieck, 1998). This parallel with our results suggests that the early visual system may have extracted a simple yet potent set of features while subject to genomic bottleneck-like constraint.

### Reinforcement learning

The results described so far demonstrate the efficacy of genomic compression in the context of supervised learning. However, supervised learning is unlikely to play a major role in animal behavior (Zador, 2019). We therefore turned our attention to reinforcement learning paradigms, in which an agent seeks to maximize its reward in a given environment by taking actions based on its current state and its history of actions and rewards. The actions are determined by a policy which maps the agent’s state to actions. Learning in this context consists of adapting the policy. Many of the most successful modern approaches use ANNs to implement the policies (Silver et al., 2016).

We first tested the genomic compression algorithm on the ANN-based policies used for solving Beam Rider (Bellemare et al., 2013), a video game (Fig. 4). In this task, the input is a set of 80 × 80 pixels and the output is one of 9 actions (e.g. move left, fire, etc). We used the dueling deep Q learning algorithm (Wang et al., 2016) to train a standard p-network with 3.3 × 10^6^ parameters. With training, performance typically doubled after several hundred episodes (Fig 4C). We then compressed the p-network 492-fold using a g-network with about 10^4^ parameters. Initial performance of the compressed network was nearly asymptotic with little or no training, indicating that the g-network was able capture almost all of the structure inherent in the connections of the p-net. We then tested greater levels of compression, and found that the innate performance of the compressed network remained excellent up to about 3500-fold compression (Fig 4D). Similar results were obtained for another video game, Space Invaders (see Supp Fig. S.5). These results show that the genomic compression approach is not limited to supervised learning, but can be readily extended to a reinforcement learning setting.

**Figure 4:**
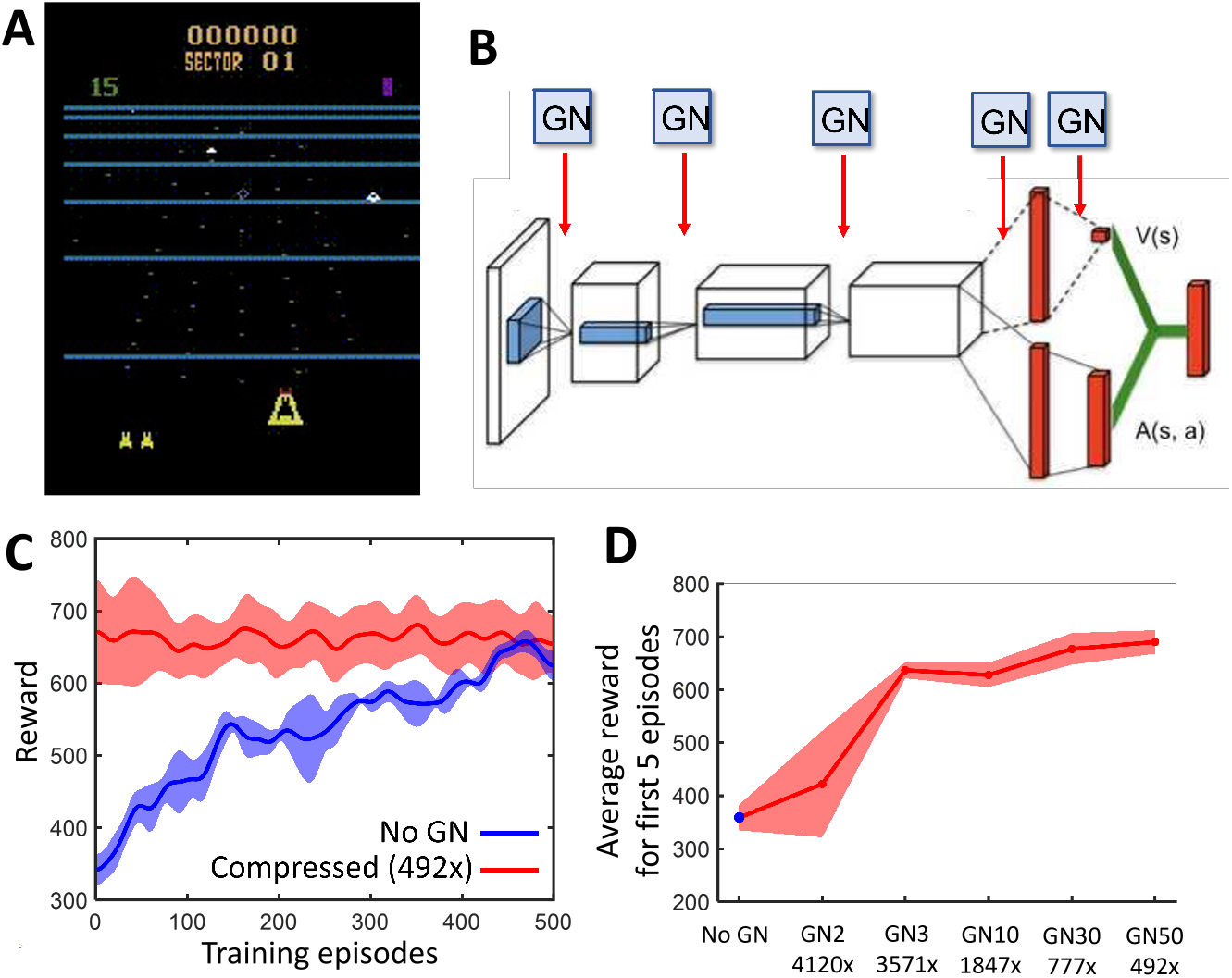
Genomic bottleneck approach to reinforcement learning. (A) Application to the video game Beam Rider, in which the goal is to shoot down enemy ships. (B) Network architecture of p-net, which consisted of 5 layers, each of which was compressed by a corresponding g-net. (C) Dynamics of learning for uncompressed network (*blue*) is slower than for a 492-fold compressed network (*red*), which achieves nearly perfect performance on episode 0. Shaded regions show standard deviation of the data. (D) Performance over first 5 episodes as a function of compression. Nearly perfect initial performance is achieved by the x3570-fold compression.

We next tested the genomic compression algorithm on a more challenging reinforcement learning task, the “Half Cheetah” (Fig. 5). In this task, a simulated cheetah must learn to maximize its forward velocity in a simulated physics environment, Mujoco (Todorov et al., 2012). Here, the state space consists of the positions, angles and velocities of 8 joints, and the continuous action space consists of the forces applied to those joints. With a random initialization, the cheetah cannot move forward (Fig. 5A, *top*). After several thousand episodes of training, the cheetah sometimes learns to move forward in a conventional way, but typically adopts unconventional solutions such as tumbling on its head or flipping upside down and gliding along its back (Fig. 5A, *middle*), a phenomenon sometimes referred to as “reward hacking”. Such solutions often yield rewards comparable to those obtained by conventional solutions, and can be viewed as a form of overfitting.

**Figure 5:**
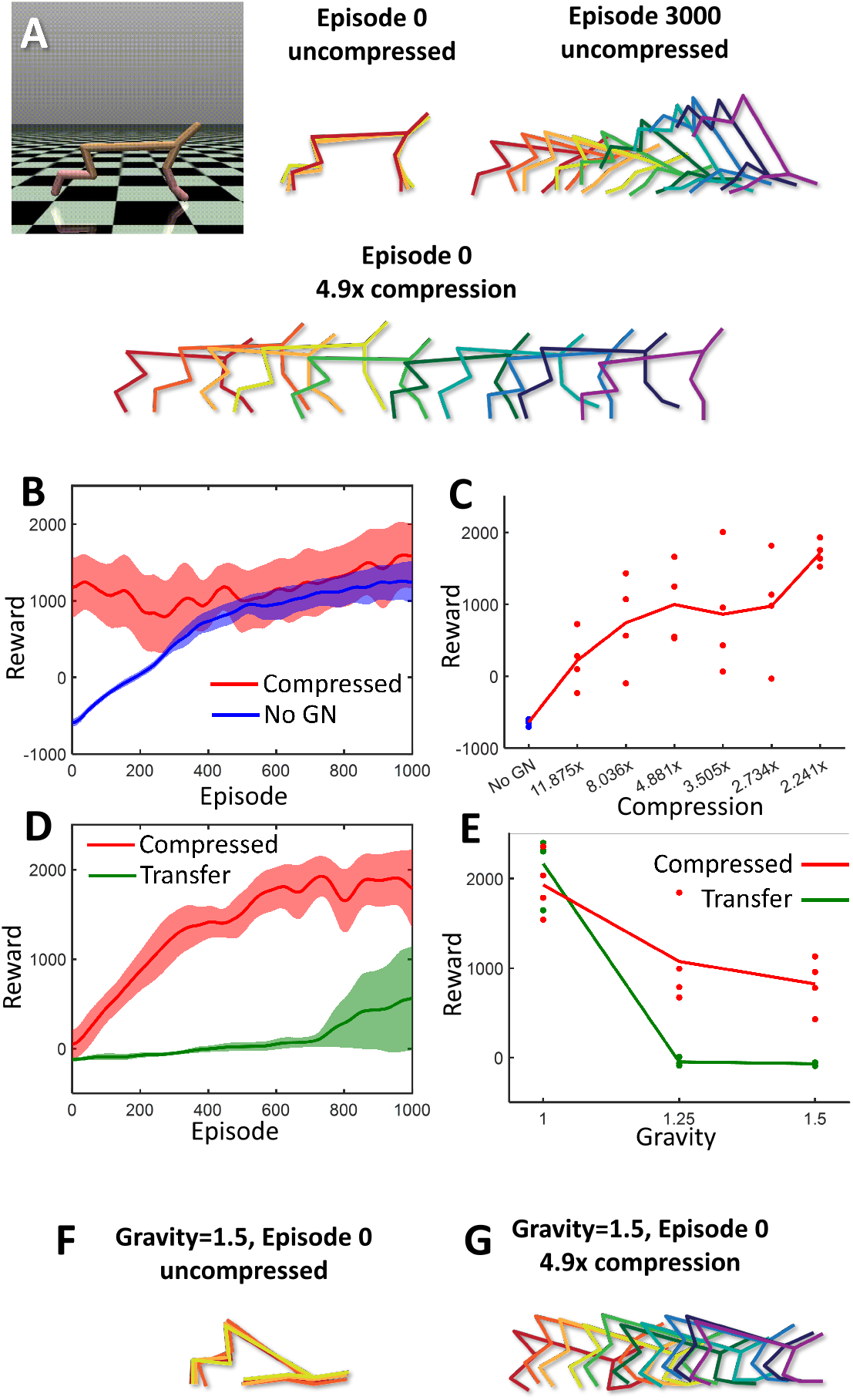
Genomic bottleneck applied to Half Cheetah. (A) Snapshots of sample episodes. (*top, middle*) Upon initialization without compression, the Half Cheetah typically flails about, largely in place. (*top, right*) After 3000 episodes of training, the agent finds an effective but often unconventional strategy for moving forward. In this example, the strategy involves intermittent sliding on its chin. (*bottom*) The agent trained with the genomic bottleneck approach adopts an effective and conventional strategy even on episode 0. (B) Learning time course for x4.9-fold compressed (*red*) vs. randomly initialized (*blue*) networks. Shaded region shows standard deviation of the data for four independent training runs. (C) Initial performance of the Half Cheetah network averaged over the first 5 episodes as a function of compression. Performance is shown for individual genomic networks (dots) and the average (solid line). (D) The results of weight transfer to the environment with higher gravity. Learning time course for x4.9-fold compressed (*red*) vs. uncompressed (1x, direct transfer) (*green*) networks. (E) Performance averaged over the first 400 training episodes as a function of gravity scale. Dots/lines show results for individual networks and their average. Colors are the same as in (D). (F,G) Skeleton diagrams showing subsequent time steps for networks initialized by direct transfer (F) and 4.9x compressed g-net (G) in the increased gravity environment.

The genomic bottleneck algorithm was able to achieve excellent compression on this task. After 4.9-fold compression, performance on episode 0 is excellent, approaching asymptotic performance (Fig. 5B). Initial performance on episode 0 declined only modestly with increasing compression (Fig. 5C). Thus, the genomic compression algorithm could be effectively applied even in the more challenging setting of the Half Cheetah.

Interestingly, solutions obtained from p-nets initialized by g-nets did not appear to engage in reward hacking, unlike those obtained following random initialization; following compression, only conventional strategies without tumbling and flipping were learned (Fig. 5A, *bottom*). This suggested that compression was acting as a regularizer, discouraging overfitting by reward hacking. To further explore this, we tested performance in a modified environment in which gravity has been increased by 50%. (In the Mujoco simulation environment, gravity can be conveniently modified with a single parameter, which can be viewed as a surrogate for changes in body size that would occur over an animal’s lifetime). In this modified environment, initial performance at episode 0 even for the p-network initialized by a g-network is poor, but performance improved relatively quickly after several hundred trials (Fig. 5C, *red*). By contrast, conventional transfer learning, in which the fully trained uncompressed network is used in the new environment, learned much more slowly (Fig. 5D, *green*; Fig. 5E). The superior transfer learning observed with the genomic bottleneck approach arises because the reward-hacked strategies observed in networks trained without compression do not generalize well to the new environment, so the agent must first unlearn these unconventional strategies. Taken together, our results suggest that the genomic bottleneck approach can be effectively applied to both supervised and reinforcement learning problems.

## Discussion

We have proposed that a genomic bottleneck arises inevitably because of the need for a relatively low-capacity genome to specify the complex neural circuits required for innate behaviors. We argue that under a wide range of conditions, there is evolutionary pressure for organisms to be born with as much innate ability as possible, and thereby to maximize their fitness at birth (Fig. 2E). This leads to a model in which genomes, and the circuits they encode, are co-optimized in nested loops: an inner loop corresponding to “learning” in animals, and an outer loop corresponding to “evolution.” Our results suggest that dividing the usual machine learning problem into such nested loops linked by a low-information bottleneck serves as a regularizer on the resulting neural circuit, guiding it to find simple circuit motifs that can be reused and can adapt with changes in the environment.

The genomic bottleneck can be viewed as a constraint forcing lossy compression of the weight matrix (Eq. 1). The idea of minimizing the description length, or Kolmogorov Complexity, of the weights has a long history (Hinton and Van Camp, 1993; Schmidhuber, 1997). Although the genomic bottleneck algorithm was motivated by considerations of the relative size of the genome and the connectome, it has close parallels with the “information bottleneck” method (Tishby et al., 2000). We hypothesize that, by squeezing the neural circuit diagram through a much smaller genome, evolution has extracted the most useful and important network motifs.

To perform tasks, the compressed (genomic) representation must be uncompressed into a functional network through a process analogous to neural development. For reasons of efficiency, we have used gradients to optimize both the inner and outer loops. Evolution, which can be viewed as a form of optimization that does not exploit a gradient, is in general a relatively slow and inefficient algorithm, successful because it operates on massive numbers of individuals in parallel over hundreds of millions of years. The feedback in our algorithm that guides the gradient from each generation—the fact that the trained weight matrix in the *k*^*th*^ generation is used to modify the genome in the (*k* + 1)^*st*^ generation—can be viewed as a form of Lamarckian evolution, and is, as such, biologically unrealistic. The net effect of our approach, however, is similar to Darwinian evolution. Our algorithm can also be seen as an implementation of the Baldwin effect (Baldwin, 1896; Hinton and Nowlan, 1987), according to which, if the ability to learn a particular behavior rapidly conferred a selective advantage, that ability would, over the course of evolution, be “genetically assimilated” and might appear to have arisen through a Lamarckian process.

We developed a genomic bottleneck algorithm that could achieve several orders of magnitude compression on standard supervised and reinforcement learning benchmarks. Although it might seem surprising that these networks could be so highly compressed with relatively little loss of performance, our results are consistent with a considerable literature on network compression (Choudhary et al., 2020). For example, a standard technique—weight pruning—can eliminate 90% of parameters with minimal loss of accuracy (LeCun et al., 1990; Han et al., 2015). Network pruning can be viewed as a method of network compression, complementary to the genomic bottleneck mechanism considered here. Convolutional neural networks (LeCun and Bengio, 1995) represent an example of network pruning whereby each neuron only connects to a small fraction of other neurons in lower layers. Another example is provided by the lottery ticket hypothesis, according to which the number of weights in a well-performing network can be substantially reduced by discovering “winning ticket” sparse subnetworks (Frankle and Carbin, 2018). Cortical networks are inherently sparse, with each neuron connecting to only a minute fraction of other cortical cells. Evolution selects sparse and functionally important connections due to physical constraints, such as a space and time limitations (Chklovskii et al., 2002). Even after sparsification, the organization of cortical connections cannot be encoded in the genome with the single neuron precision. Thus, pruned connectivity is a default solution to the evolution of cortical fitness, and does not by itself resolve the discrepancy between cortical and genomic information capacities; even sparse connectivity must be further compressed through the genomic bottleneck. Here we study the additional rules that can encode both fully and sparsely connected networks.

Our results contribute to a growing literature highlighting the importance of inductive biases in machine learning. Much of this literature is focused on achieving faster learning. Perhaps the most successful examples are convolutional neural networks (LeCun and Bengio, 1995), which exploit the translational invariance of images with an architecture inspired by the structure of receptive fields in early sensory cortex (Hubel and Wiesel, 1962). However, the present work—inspired by evolutionary constraints (Fig. 2D)—is focused not on faster learning but rather enhanced initial performance, a goal that has received comparatively less attention (but see (Gaier and Ha, 2019)). Indeed, in our experiments we find that genome-initialized networks start off at a higher level of performance but then follow the same trajectory as randomly-initialized networks (Fig. 3D). Although these results highlight the potential dissociation of two distinct process—better initial performance and faster learning through inductive biases—there is likely strong evolutionary pressure to maximize both.

The relative importance of genomically-encoded innate structures in determining specific human abilities such as language has been hotly debated, but the importance of innate factors to the behavior of other animals is less controversial. For both humans and other animals, the better question is usually not whether a behavior is innate, but rather how innate and learned factors interact. For example, the propensity to form “place fields” in the hippocampus is innate—a map of space emerges when young rat pups explore an open environment outside the nest for the very first time (Langston et al., 2010)—but the content of place fields is learned, as new place fields form whenever the animal enters a new environment. In this example it appears that, as suggested by our experiments, innate performance has been maximized by providing a scaffolding for place fields to appear.

The genomic bottleneck takes its inspiration from fundamental considerations about the evolution and development of brain circuits. Although the genomic bottleneck algorithm builds on existing machine learning techniques, and yields surprisingly effective performance, we have not attempted to optimize this approach to compete with state-of-the-art benchmarks. The bottleneck framework is potentially quite rich, and could be extended in several directions. For example, we have not explored variations in the structure of the genomic network, *e*.*g*. by imposing a sparseness constraint. Such a constraint would have the physical interpretation of limiting interactions among surface neuronal markers. Similarly, at present, each layer is compressed with its own genome, but it would be natural to attempt to extract regularities among layers by encoding them with a single genome. Furthermore, in the current formulation the decoding of the genomic network is deterministic, whereas developmental rules are often stochastic, so the decoding framework might be generalized to include rules like “let each neuron connect on average to 10% of nearby neighbors” or a more complex stochastic rule (Stöckl et al., 2021). Finally, the framework could be extended to co-optimize the learning rules and the wiring.

## Acknowledgments

We are grateful for many helpful comments from Mike DeWeese, Saket Navlakha, Blake Richards, Tatiana Engel, Xaq Pitkow, Kaleb Vinehout, Barak Pearlmutter. We would like to acknowledge support from Deep Valley Labs.

## Methods

### MNIST/F-MNIST dataset

For MNIST and F-MNIST datasets, we used a fully connected 2-layer network that included 800 hidden layer ReLU units (Simard et al., 2003). We did not use data augmentation for simplicity and our network could be trained to 98% performance on testing data. The number of parameters in the MNIST network was therefore 28^2^ × 800 + 800 × 10 weights and 800 + 10 biases amounting to the total of 636010. We used three g-networks to encoded two weight matrices and one bias vector for the hidden layer. The ten biases for the output layer were not compressed since the corresponding g-network would include more than ten parameters. The structures for various configurations of the three g-networks are listed in Table 1. The schematic of the structure of g-network for MNIST dataset is shown in Fig. S.1A.

**Table 1:** The structure of g-network for MNIST dataset

| g-network name | g-network structure | Bias net structure | g-network parameters | MNIST parameters | Compression |
| --- | --- | --- | --- | --- | --- |
| GN5 | $(20 - 10 - 5 - 1) \times 2$ | | 613 | | 1038x |
| GN20 | $(20 - 20 - 10 - 1) \times 2$ | 10 - 5 - 1 | 1,353 | 636,010 | 470x |
| GN30 | $(20 - 30 - 10 - 1) \times 2$ | | 1,973 | | 322x |

Each neuron was described by a binary label of length 10. For the neurons in the input image, the label encoded two coordinates of the neuron’s position in the image, 5 bits for the “X” and “Y” coordinates. Both coordinates were represented by the Gray code. The neurons in the hidden layer were represented by simple binary codes ranging from 1 to 800, since their order is of no particular importance. The ten neurons in the output layer were encoded by the one-hot vector of 10 bits. Each neuron in the networks was therefore described by a ten-bit label. The inputs into each of the two g-networks that generated weight matrices represented pairs of neurons and had the length of 20 bits. The output of g-network is the value of the corresponding weight between two input neurons and was a single real number (Table 1). For the network generating biases for the hidden layer, the input contained the binary label for the neuron and was therefore 10 bits long.

There are several options to train g-networks. The simplest one is to use end-to-end backpropagation from the dataset (MNIST) to g-network using automatic differentiation of the deep learning library (PyTorch, Fig. S.1A). We found this implementation to be inefficient as it involves generation of the entire set of weight matrices for each mini-batch of MNIST images. Instead, we developed the intermittent training paradigm. Before the first iteration, the g-networks are initialized randomly. On each iteration of this method, we start with the g-networks generating weights of the MNIST network (p-net). We then train the p-network using a subset of images. For the MNIST network, this training used 10,000 images from the training set or 1/6th of the epoch. This yielded a higher performance p-network with the set of weights described by matrix *W*_*n*_. We then train g-networks to approximate this weight matrix by backpropagating the difference between the g-network output 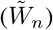 and *W*_*n*_. We used different number of weights to train each of the three g-networks on each iteration (10^5^, 10^4^, and 10^4^ for hidden layer, output layer, and hidden layer biases g-networks, respectively). This amounted to about 1/6th of all weights and further accelerated training in each generation. We then used the adjusted g-networks to generate the complete set of p-network weights 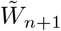 that served as initial conditions for the next generation (Fig. S.1B). This set of iteration mimicked real biological evolution as it alternated the generation of p-nets from g-networks, analogous to the neural development, and improvement of p-nets similar to the natural selection. We repeated these iterations 500 times to achieve the asymptotic performance.

### CIFAR10/SVHN datasets

In this example we used all convolutional 9-layer implementation of network (Springenberg et al., 2015) (Fig. S.1B). Between layers 3 and 4, we included the dropout layer with 50% dropout probability. The network could be trained to 89% correct rate without data augmentation. Each layer in this 9-layer CNN was generated via two g-networks, one for the weight matrix and one for the biases. To provide inputs into the g-networks for weights, we described positions in the weigh matrix by a 20-bit binary number. For the lowest 8 layers, the binary number was composed of 2+2 bits containing Gray code representation for the input coordinates within the CNN kernel, 8 bits representing the input filter type, and 8 bits representing the output filter type. The latter two components were formed as consecutive binary numbers, since input/output filter identity is not expected to form a continuous topographic space. In this representation, the input neurons were identified by a 12-bit binary label (2 + 2 Gray + 8 binary) while the output neurons are identified by the 8-bit label. The binary labels for the last CNN layer were composed of 1+1 binary Gray (dummy) bits representing two coordinates within the kernel, a 8-bit binary number representing the input filter, and a 10-bit one-hot vector encoding the output class. The bias networks for the first eight layers received an 8-bit label encoding the output filter. The bias network for the last layer received the one-hot vector encoding one of the ten output classes. The structures of different g-networks used are summarized in Table 2.

**Table 2:** The structure of g-network for CIFAR10 dataset

| g-network name | g-network structure | Bias net structure | g-network parameters | CIFAR10 parameters | Compression |
| --- | --- | --- | --- | --- | --- |
| GN10 | $(20 - 10 - 10 - 1) \times 9$ | | 3,798 | | 361x |
| GN30 | $(20 - 30 - 10 - 1) \times 9$ | 8 - 10 - 1 | 9,378 | 1,369,738 | 146x |
| GN50 | $(20 - 50 - 10 - 1) \times 9$ | | 14,958 | | 92x |

To train the 18 g-networks described above, we used the intermittent training strategy described for the MNIST network. We used minibatch sizes of 10, 100, and 1000 for bias g-networks for layers 1-3, 9, bias g-networks 4-8, and all weight matrix g-networks, respectively. We used the minibatch size of 128 to train the CIFAR10 network. We used SGD optimizer for CIFAR network with the learning rate of 0.05 and momentum of 0.9 for stability. We used Adam optimizer for all g-networks. In each iteration, we first trained CIFAR10 network using 10 complete epochs, i.e. 10 complete sweeps through the entire training set of images. In the second step, we trained weight and bias g-networks using 2 and 10 epochs respectively (we trained g-network weight networks for layer 1 and 9 using 12 epochs in each iteration) to match the CIFAR network adjusted weights resulting from the first step. This sequence of two steps was repeated 500 times.

Because our network was relatively deep (9 layers), we encountered a problem with initialization of g-networks. Indeed, if g-networks are initialized randomly, they produce p-nets that are far from the optimal fixed point. We found that such p-nets are impossible to train. This problem is exacerbated in moderately deep p-nets due to the exponential divergence of initialization errors from layer to layer. In practice, such p-nets return zero activations, which yield no gradients of weights. To solve this problem, we implemented the weight annealing strategy. In each iteration of our algorithm (out of 500), before the p-network was trained, the weight matrices and biases of the p-network were combined from the results of CIFAR training in the previous iteration, *W*_*n*−1_, and the weights generated by g-network in the previous iteration, 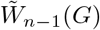:

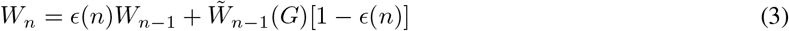

The coefficient *ϵ*(*n*) determined the degree to which the inputs from g-networks affect the p-net’s weight matrix. If *ϵ*(*n*) = 1, the weight matrix of p-network is entirely determined by the result of previous iteration of CIFAR training and is not sensitive to the inputs from g-net. In the other extreme, when *ϵ*(*n*) = 0, the values of p-network weights and biases are entirely generated by the g-networks. We therefore assumed that *ϵ*(*n*) ≈ 1 in the beginning of training, when g-networks are naive, and *ϵ*(*n*)→ 0 in the end of training. We adopted an exponential annealing schedule with *ϵ*(*n*) = exp(−*n*/*λ*), where parameter *λ* = 20 determined the number of iterations in the intermittent training over which the g-networks are assumed to be naive and irrelevant, and initialization using g-networks is assumed to be detrimental. Since, in our approach, the total number of iterations is 500, the initialization period is negligibly short compared to the whole training (*λ* « 500).

### RL Methods

We performed experiments on two reinforcement learning tasks from t0068e OpenAI environment (Brockman et al., 2016): BeamRider and Half Cheetah. The details of the experiments follows.

For the first experiment, we tested the genomic bottleneck on solving the BeamRider task, which is part of the Atari benchmark (Bellemare et al., 2013). The task in the beam rider is to traverse a series of sectors where each sector contains 15 enemies and a boss at the end. Additionally, the agent needs to avoid or destroy the debris coming its way. The agent is equipped with three torpedoes that can be used to kill the enemies or destroy the debris.

We used the pixel images of Atari frames as our state space. Atari frames are 210 × 160-pixel images with a 128 color palette. This makes it computationally expensive to use it directly as input to the network. In order to reduce the dimension of the input, we performed the following preprocessing steps on the images: conversion of the RGB image to gray scale, cropping it to get an image of size 190 × 144 and finally resizing it to the size 84 × 84. The action space is of size 9 with various possible actions: fire up, fire, up, left, right, left fire, right fire, up left, up right, and do nothing. The score obtained while playing the game was used as the reward signal.

**Figure S.1:**
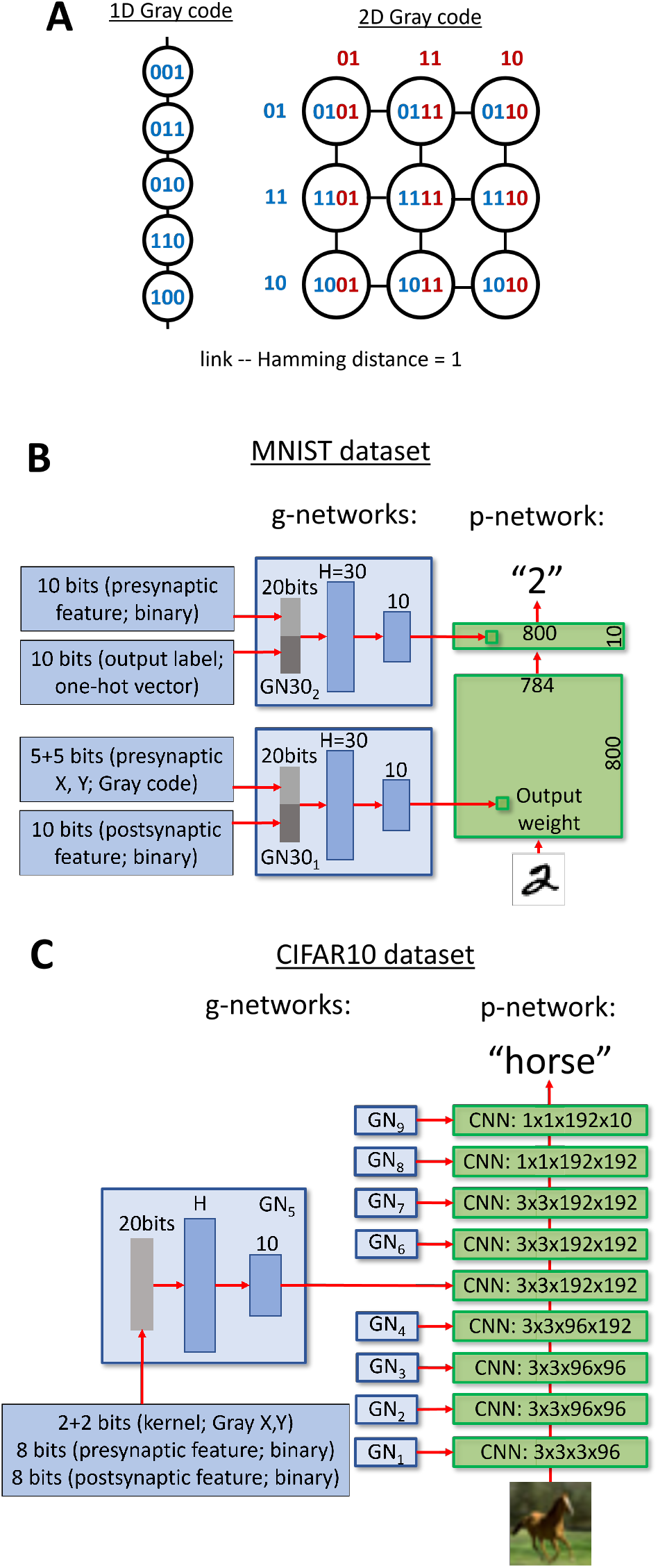
Networks’ structure for MNIST and CIFAR datasets (A) Neuron labels are assigned by a 2D Gray code, so that neurons close in image space have similar (Hamming distance = 1) labels. The structure of g-networks for MNIST (B) and CIFAR10 (C) datasets.

**Figure S.2:**
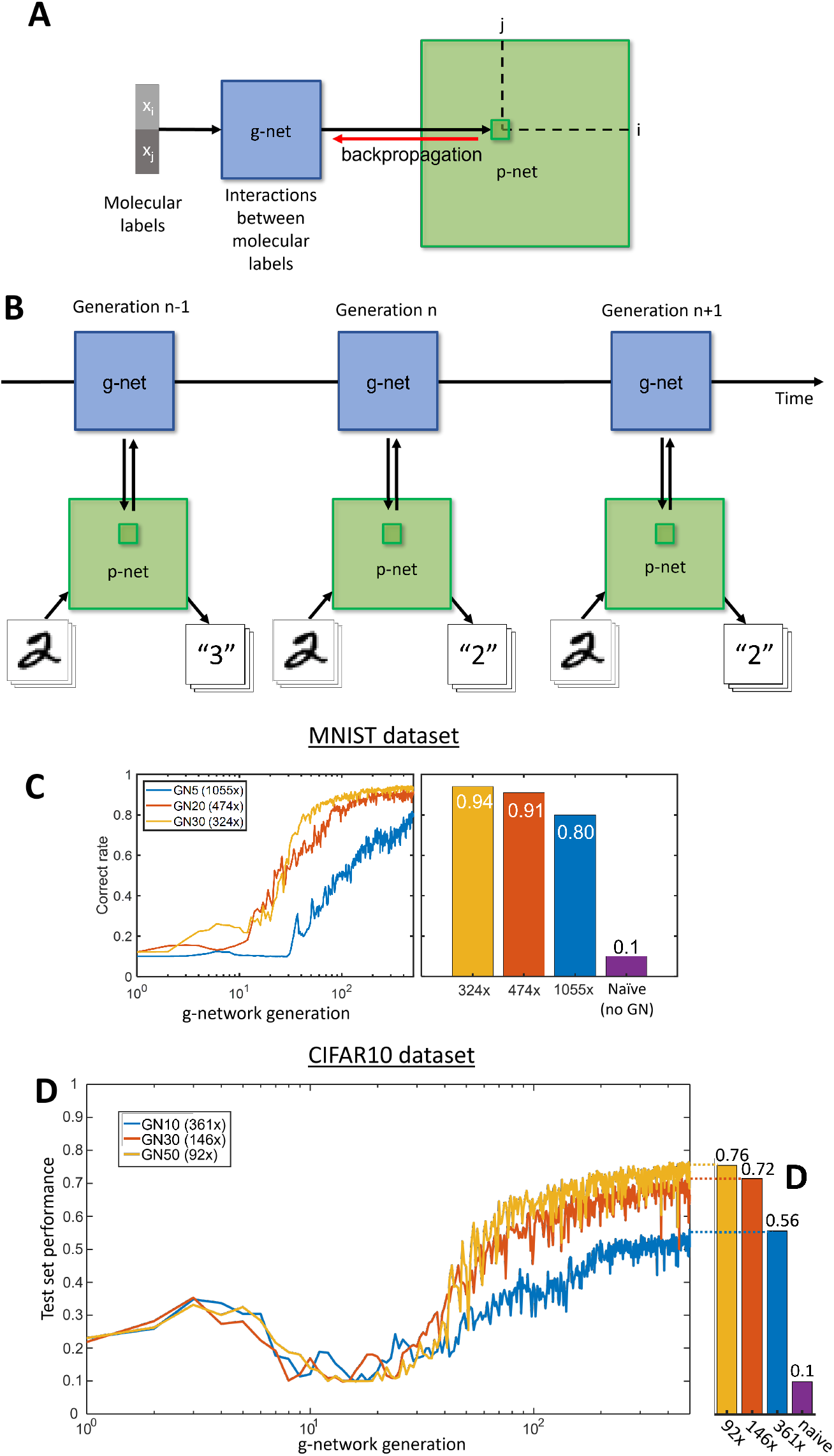
Training strategies of g-networks. (A) End-to-end backpropagation of errors from the output of the p-network to g-networks. This implementation is slow since it involves generating a complete set of p-network weights and biases for each p-network minibatch. (B) The intermittent training strategy. G-network of generation n-1 is used to generate the p-network (down arrow, generation n). The p-network is trained using several minibatches without backpropagation of the gradients into g-network. Then, the g-network is trained to match the adjusted p-network (up arrow in generation n). The resulting g-network in generation n is used to generate the p-networks in the next step (n+1). (C) The dynamics of training of g-network for the MNIST dataset. (D) The dynamics of training for CIFAR10 dataset. The small bump in performance at generations 2-7 is due to the annealing strategy used to initialize g-networks in this case [Eq. (3)].

**Figure S.3:**
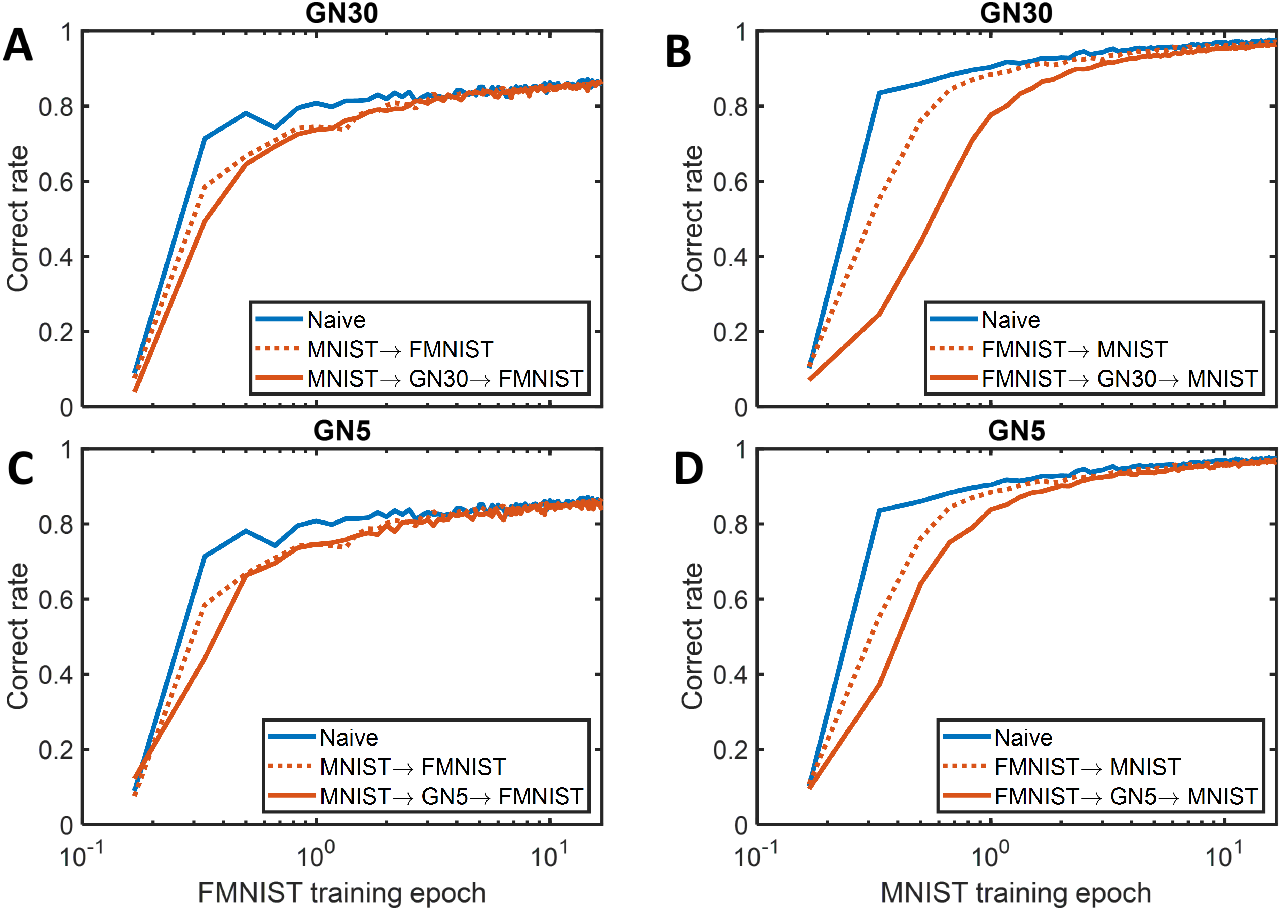
The results of weight transfer between MNIST and FMNIST datasets compared to training networks based on random initialization (blue). Weight transfers using the entire set of weight matrices (dotted red lines) are contrasted to the transfer using g-nets (solid red). Two types of g-nets are used, with low (A, B) and high (C, D) compression. Transfers from MNIST to FMNIST (A, C) and from FMINST to MNIST (B, D) show qualitatively the same results – training from scratch is better for this pair of datasets, even if the transfer occurs using the entire weight matrix (dotted, no compression).

**Figure S.4:**
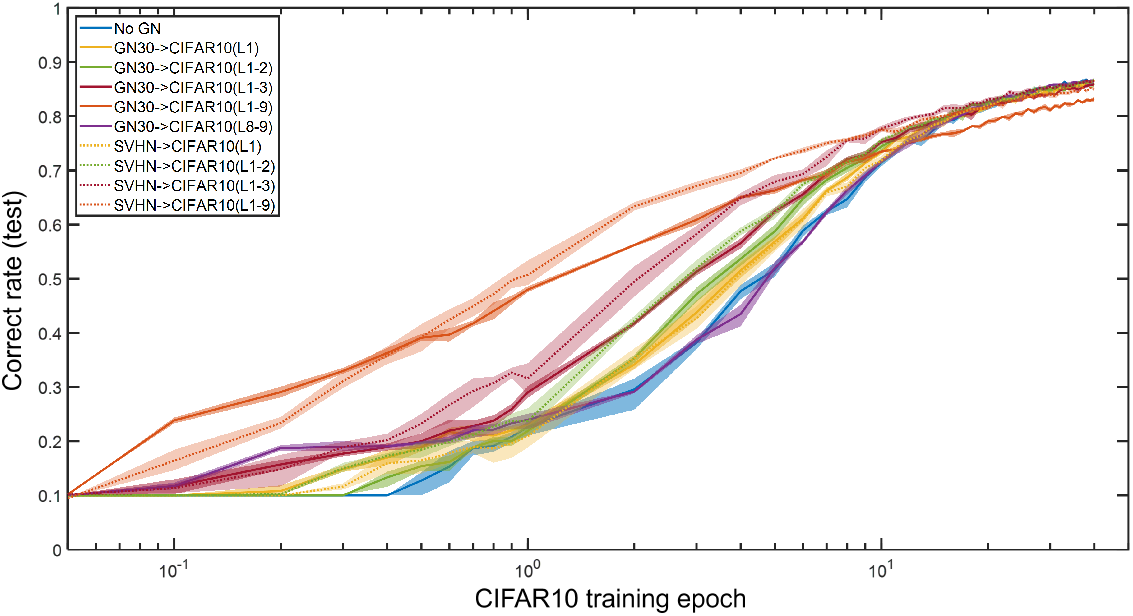
The results of reverse transfer experiments from SVHN to CIFAR10 dataset. Training of a naïve weight matrix from scratch (blue) is contrasted with direct weight transfer (dotted lines) and transfer using the g-net (solid lines, GN30). Different sets of layers were transferred as indicated by the color map. Overall, these results show that, although different scenarios of weight transfer yield faster training than training from scratch (blue), the difference between uncompressed (dotted) and g-net compressed (solid) cases is not significant.

For the p-network, we used Dueling Deep Q Network (DDQN) (Wang et al., 2016) to train the on this task. Dueling DQN breaks the calculation of Q value into two parts. *Q*(*s, a*) = *V* (*s*) + *A*(*s, a*), where *V* (*s*) represents the Value of state “s” and A represents the Advantage of performing action “a” while in state “s.” The value of a state is independent of action. This aids us in avoiding the overshoot of the Q-value that occurs in vanilla deep Q networks (Mnih et al., 2015). Since the states are independent of action, action would not have a high Q-value to train on and thus Q-value would not overshoot. This makes the training smooth and less time-intensive

The p-network architecture includes three 2-D convolutional layers and three fully-connected layers with a total size of ∼3M parameters. The input to the DDQN is our pre-processed image of size 84 × 84 and the output is the Q value associated with each action.

For the second task, we moved to a more challenging task i.e. ‘Half Cheetah’ (Todorov et al., 2012). One challenging aspect of this task is that the Half Cheetah uses continuous action spaces rather than discrete unlike most of the Atari games. This leads to a performance deficit in value approximation-based methods like DDQN.

The state-space of the Half Cheetah is of size 17 which contains the position, angles, and velocities of 8 joints (6 hinge + 2 slider joints). The action space is a 6-dimensional continuous space consisting of the torques applied to each hinge joint respectively. Since we want to maximize the velocity with minimum force generation, the reward function needs a combination of two components. The first component provides a reward for velocity and the second component gives a penalty for using more force.

For training the p-network for this task, we need a training method that can deal with continuous action spaces. The class of methods that addresses this domain includes policy gradient methods because they can directly approximate the policy of the agent from the state. Proximal policy optimization (PPO) (Schulman et al., 2017) is one of the most popular policy gradient methods and is heavily used for such continuous action spaces tasks. One of the main reasons behind this is that policy gradient methods have a convergence problem which is usually addressed by the natural policy gradient. However, in practice, natural policy gradient involves a second-order derivative matrix which makes it not scalable for large-scale problems. PPO uses a slightly different approach. Instead of imposing a hard constraint, it formalizes the constraint as a penalty in the objective function. By not avoiding the constraint at all costs, PPP was able to use a first-order optimizer like the Gradient Descent method to optimize the objective resulting in faster convergence. The p-network architecture of the Half Cheetah consists of three fully connected layers with 6092 trainable parameters. The input to our p-network is the state vector of the environment and the output is the action the agent should take given that state.

**Figure S.5:**
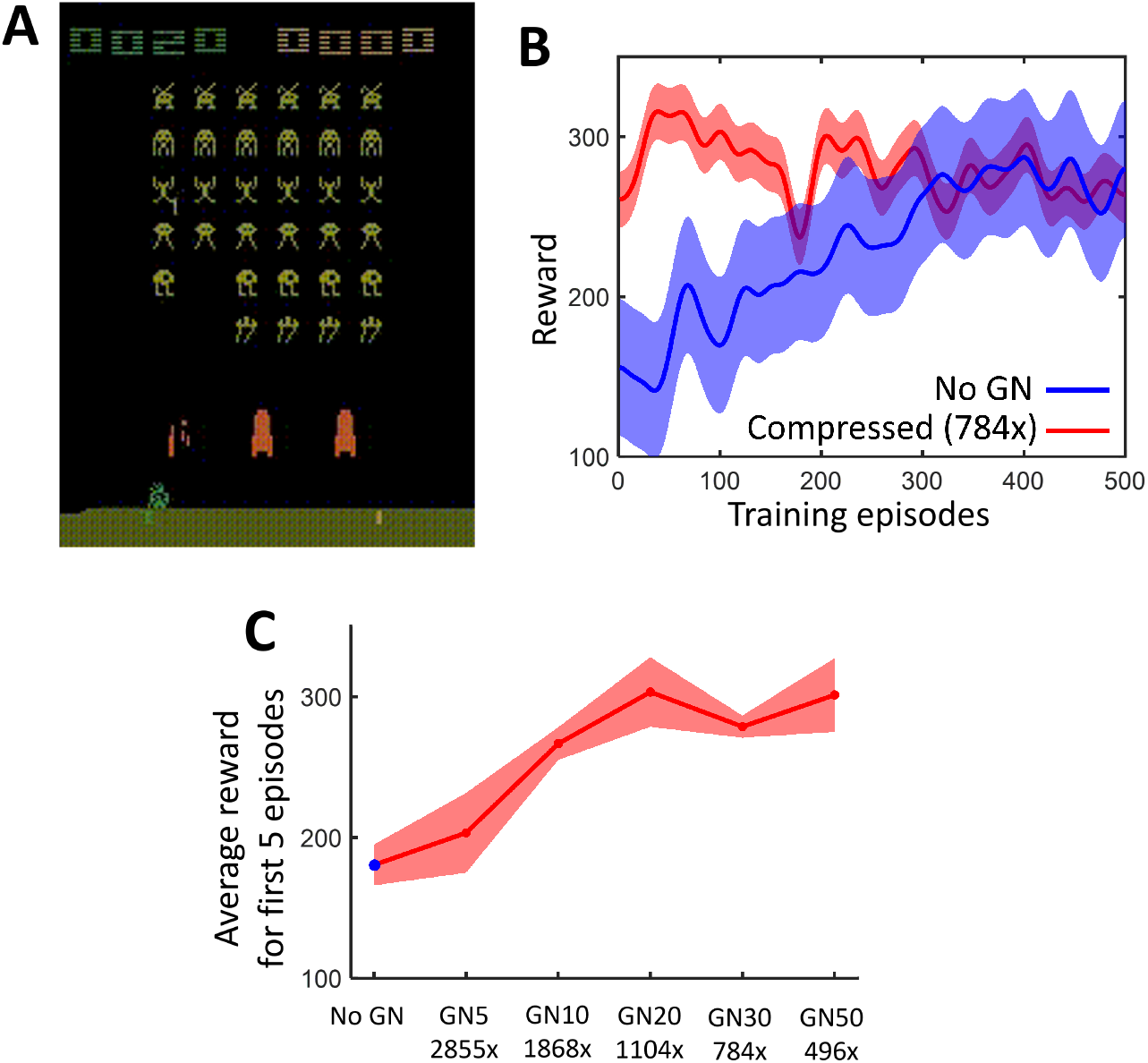
Application of genomic bottleneck to the Atari video game Space Invaders. (A) Screen shot of the game. We used the same architecture as in the Beam Rider case. (B) Dynamics of learning for uncompressed network (*blue*) is slower than for a 784-fold compressed network (*red*), which achieves nearly perfect performance on episode 0. Shaded regions show standard deviation of the data. (C) Performance over first 5 episodes as a function of compression.

